# Targeting IDH1 as a pro-senescent therapy in high-grade serous ovarian cancer

**DOI:** 10.1101/472613

**Authors:** Erika S. Dahl, Raquel Buj, Kelly E. Leon, Jordan M. Newell, Benjamin G. Bitler, Nathaniel W. Snyder, Katherine M. Aird

**Author notes:** Corresponding Author: Katherine M. Aird, Assistant Professor, Department of Cellular & Molecular Physiology, Penn State College of Medicine, Hershey, PA.

## Abstract

Epithelial ovarian cancer (EOC) is the deadliest gynecological cancer. High-grade serous carcinoma (HGSC) is the most frequently diagnosed and lethal histosubtype of EOC. A significant proportion of HGSC patients relapse with chemoresistant disease. Therefore, there is an urgent need for novel therapeutic strategies for HGSC. Metabolic reprogramming is a hallmark of cancer cells, and targeting metabolism for cancer therapy may be beneficial. Here we found that in comparison to normal fallopian tube epithelial cells, HGSC cells preferentially utilize glucose in the TCA cycle and not for aerobic glycolysis. This correlated with universally increased TCA cycle enzyme expression in HGSC cells under adherent conditions. To further differentiate the necessity of TCA cycle enzymes in ovarian cancer progression, we found that only wildtype isocitrate dehydrogenase I (IDH1) is both significantly increased in HGSC cells in spheroid conditions and is associated with reduced progression-free survival. IDH1 protein expression is also increased in primary HGSC patient tumors. Pharmacological inhibition or knockdown of IDH1 decreased proliferation of multiple HGSC cell lines by inducing senescence. Mechanistically, suppression of IDH1 increased the repressive histone mark H3K9me2 at proliferation promoting gene loci (*PCNA* and *MCM3*), which led to decreased mRNA expression. Altogether, these data suggest that increased IDH1 activity is an important metabolic adaptation in HGSC and that targeting wildtype IDH1 in HGSC alters the repressive histone epigenetic landscape to induce senescence. Therefore, inhibition of IDH1 may act as a novel therapeutic approach to alter both the metabolism and epigenetics of HGSC as a pro-senescent therapy.

## Introduction

Among all gynecological cancers, epithelial ovarian cancer (EOC) is the most lethal due to dissemination into the peritoneal cavity and omental seeding in late stage disease (1). High-grade serous carcinoma (HGSC) is the most common histosubtype of EOC and has the worst prognosis (2). The five-year survival rate for HGSC patients is approximately 47% due to limited screening options and late stage diagnosis (3). This percentage decreases to 20-35% in women diagnosed with stage III and IV HGSC. Due to their characteristic *TP53* mutations, many HGSC patients initially respond to standard-of-care platinum and taxol chemotherapies; however, a significant portion relapse with chemoresistant disease (4, 5). PARP inhibitors were recently approved as a maintenance therapy for ovarian cancer (6, 7). Although these inhibitors show promise, especially for homologous recombination-deficient ovarian cancer (8, 9), some patients do not respond and resistance to these drugs has recently become evident (10, 11). Therefore, there is an urgent need for novel therapeutic strategies for HGSC patients.

Cellular senescence is a state of stable cell cycle arrest that is induced by multiple stimuli, including shortened telomeres, oncogene activation, DNA damage, and certain therapeutics (12). Senescent cells are characterized by many hallmarks such as increased β-galactosidase activity (termed senescence-associated β-galactosidase or SA-β-gal), increased repressive histone marks, such as repressive di- and tri-methylation marks of histone 3 lysine 9 (H3K9me2/3) at proliferation-promoting genes (termed senescence-associated heterochromatin foci, SAHF), decreased incorporation of the thymine analog bromodeoxyuridine (BrdU), and decreased proliferation (13–15). Due to the decrease in proliferation, therapy-induced senescence is considered a desirable therapeutic outcome (16–18). Interestingly, recent evidence suggests that therapy-induced senescence leads to a better 5-year survival rate for HGSC patients (19).

Tumor cells are characterized by metabolic reprogramming in order to maintain uncontrolled cell proliferation and growth (20). Tumors specifically reprogram metabolic pathways to allow for the increased need for biomass, such as nucleotides, lipids, and other macromolecules (21, 22). This metabolic reprogramming is increasingly thought to modulate epigenetic marks, including histone and DNA methylation and histone acetylation (23). Altered metabolism in cancer cells provides a unique opportunity to exploit these changes as a targeted therapy (20). Cancer cells undergo anaerobic glycolysis and generate lactate, even in the presence of oxygen (24), termed the Warburg Effect. Although the Warburg effect is a common feature of many cancers, TCA cycle metabolism remains critical for both ATP production and macromolecule synthesis (25). Indeed, recent evidence suggests that multiple cancer types have increased TCA cycle metabolism(26), although less is known about TCA cycle activity in HGSC. Therefore, targeting the TCA cycle in cancer may serve as a novel therapeutic strategy.

Isocitrate dehydrogenase I (IDH1) catalyzes the oxidative decarboxylation of isocitrate to alpha-ketoglutarate (αKG) in a reversible reaction (27). IDH1 is well-known for its mutation and the resulting production of the oncometabolite D-2-hydroxyglutarate (D-2HG, aka (R)-2HG) in secondary glioblastoma and acute myeloid leukemia (AML) (28, 29). However, recent work demonstrates that overexpression of wildtype IDH1 also promotes cancer progression in primary glioblastoma (30). IDH1 regulates histone marks through production of αKG (30, 31), which is a co-factor for various histone demethylases (32, 33). Increased repressive histone methylation is a hallmark of senescence, and histone demethylases have been implicated in altering the epigenome of senescent cells (34–36). Therefore, depleting αKG pools by suppressing IDH1 may increase repressive histone methylation to induce senescence. While mutant IDH1 is well-characterized in several cancers (28, 29), the role of wildtype IDH1 in metabolism and epigenetics has never been investigated in the context of HGSC.

In this study, we determined that HGSC cells preferentially utilize glucose in the TCA cycle. While most TCA cycle enzymes were upregulated in HGSC cells compared to the cell-of-origin fallopian tube (37, 38), only wildtype IDH1 expression was both increased in HGSC cell lines under spheroid conditions and associated with decreased progression-free survival. Functionally, knockdown of IDH1 induced senescence by increasing repressive histone methylation at two proliferation-promoting gene loci: *PCNA* and *MCM3*. Together, these data indicate that targeting wildtype IDH1 in HGSC represents a novel strategy for this patient population by inducing senescence through epigenetic silencing.

## Materials and Methods

### Cells and Culture Conditions

Human HGSC Ovcar3, Ovcar10, and Ovcar5 cells lines were cultured in RPMI-1640 medium (Gibco) supplemented with 10% FBS according to ATCC. Human HGSC HeyA8 cell line was cultured in DMEM medium (Corning) supplemented with 10% FBS according to ATCC. The FT282 cell line was a kind gift from Dr. Ronny Drapkin (University of Pennsylvania). PFTE4, FT4-Tag, FT6-Tag, and FT33-Tag were a kind gift from Dr. Anna Loshkin (University of Pittsburgh). Fallopian tube cell lines were cultured in DMEM:F12 supplemented with 2% FBS and 1% penicillin/streptomycin under low oxygen conditions. HEK-293FT cells were used for lentiviral packaging and were cultured in DMEM medium (Corning) supplemented with 10% FBS according to ATCC. In order to induce spheroid formation, human HGSC Ovcar3 and Ovcar10 cells lines were cultured in 6-well ultra-low attachment (ULA) plates (Corning). Cells were treated with 1mM αKG (Sigma Aldrich) where indicated. All cell lines were cultured in Mycozap and were routinely tested for mycoplasma as described (39).

### Metabolomics

Metabolites were quantified by liquid chromatography- high resolution mass spectrometry after extraction of the cells by 80:20 methanol:water at −80 °C, sonication, centrifugation of protein at 17,000 rcf for 10 mins at 4 °C, evaporation of the supernatant to dryness under N_2_ gas and resuspended in 50 μl of 5% 5-sulfosalicylic acid for analysis. Ion pairing reversed phase liquid chromatography- mass spectrometry was conducted by modification of a previously published method on a Ultimate 3000 Binary UHPLC coupled to a Q Exactive HF mass spectrometer (40). Data were processed in Xcalibur (Thermo). Peak areas were normalized to stable isotope labeled internal standards spiked into each sample before extaction as follows; lactate, pyruvate to ^13^C_3_-pyruvate, succinate, malate to ^13^C_4_-succinate, (D/L)-2-hydroxyglutarate to ^13^C_5_-hydroxyglutarate, αKG to ^13^C_5_-αKG, fumarate to ^13^C_4_-fumarate, citrate to ^13^C_6_-citrate, acetyl-CoA, CoA, and succinyl-CoA to ^13^C_2_-acetyl-CoA.

### Patient Samples

Protein lysates of patients samples were provided by Dr. Bitler (The University of Colorado, Aurora, CO). The University of Colorado Gynecologic Tumor and Fluid Bank has an Institutional Review Board approved protocol (COMIRB #07-935) in place to collect tissue from gynecologic patients with both malignant and benign disease processes. All participants are counseled regarding the potential uses of their tissue and sign a consent form approved by the Colorado Multiple Institutional Review Board. The tissues are processed, aliquoted, and stored at −80 degrees. Benign fallopian tube tissue and primary HGSC tumors were homogenized with a Polytron Homogenizer (Brinkman Instruments) in radioimmunoprecipitation assay (RIPA) buffer (150 mM sodium chloride, Triton X-100, 0.5% sodium deoxycholate, 0.1% SDS [sodium dodecyl sulfate], 50 mM Tris, pH 8.0) supplemented with complete EDTA-free protease inhibitor cocktail (Roche), sodium fluoride (10 mM) and sodium orthovanadate (1 mM). Protein concentration was quantified using a bicinchoninic acid (BCA) protein assay (Thermo Fisher Scientific, Waltham, MA).

### Plasmids and Antibodies

pLKO.1-shIDH1 plasmids were obtained from Sigma-Aldrich. The mature sense TCRN numbers are as follows: shIDH1 #1: TRCN0000027253; shIDH1 #2: TRCN0000027249. The following antibodies were obtained from the indicated suppliers: rabbit anti-IDH1 (Cell Signaling), rabbit anti-Lamin B1 (Abcam), rabbit anti-Cyclin A (Abcam), mouse anti-Vinculin (Sigma-Aldrich), mouse anti-Beta Actin (Sigma-Aldrich), rat anti-BrdU (Abcam), mouse anti-PML (Santa Cruz Biotechnologies), mouse anti-γH2AX (EMD Millipore), rabbit anti-53BP1 (Bethyl), Fluorescein donkey anti-rat IgG (Jackson ImmunoResearch), Cy™3 donkey anti-mouse (Jackson ImmunoResearch).

### Lentivirus Infection

Lentivirus was packaged in 293FT cells using the Virapower kit from Invitrogen following the manufacturer’s instructions. Cells infected with viruses encoding the puromycin-resistance gene were selected with 3 μg/ml puromycin.

### BrdU Labeling and Immunofluorescence

For BrdU, cells on coverslips were incubated with 1μM BrdU for 30min. Cells were fixed for 10 min at room temperature in 4% paraformaldehyde. After washing the cells three times with PBS, cells were permeabilized with 0.2% Triton X-100 for 5 min and then postfixed with 1% PF + 0.01% Tween-20 for 30 min. After washing cells three times with PBS, cells were DNaseI treated for 10min (DNAseI). The DNaseI reaction was stopped using 20mM EDTA. After washing cells three times with PBS, they were blocked for 5 min with 3% BSA/PBS and then incubated in anti-BrdU primary antibody in 3% BSA/PBS (1:500) at room temperature for 1 h. Cells were washed three times and then incubated in FITC anti-Rat secondary antibody (1:1000) in 3% BSA/PBS at room temperature for 1 h. Finally, cells were incubated with 0.15 μg/ml DAPI in PBS for 1min, washed three times with PBS, mounted and sealed.

For immunofluorescence, cells on coverslips were fixed for 10 min at room temperature in 4% paraformaldehyde. After washing the cells three times with PBS, cells were permeabilized with 0.2% Triton X-100 for 5 min and then postfixed with 1% PF + 0.01% Tween-20 for 30 min. After washing cells three times with PBS cells were blocked for 5 min with 3% BSA/PBS and then incubated with the corresponding primary antibodies listed above in 3% BSA/PBS at room temperature for 1 h. Cells were washed three times and then incubated in FITC anti-Rabbit (1/2000) or Cy3 anti-mouse (1/5000) secondary antibody in 3% BSA/PBS at room temperature for 1 h. Finally, cells were incubated with 0.15 μg/ml DAPI in PBS for 1min, washed three times with PBS, mounted and sealed. All images were acquired at room temperature using a Nikon Eclipse 90i microscope with a 20x/0.17 objective (Nikon DIC N2 Plan Apo) equipped with a CoolSNAP Photometrics camera.

### SA-β-Gal

SA-β-Gal staining was performed as previously described (41). Cells were fixed for 5 min at room temperature in 2% formaldehyde/0.2% glutaraldehyde in PBS. After washing the cells twice with PBS, cells were stained at 37°C overnight in a non-CO2 incubator in staining solution (40 mM Na_2_HPO_4_, 150 mM NaCl, 2 mM MgCl_2_, 5 mM K3Fe(CN)6, 5 mM K4Fe(CN)6, 1 mg/ml X-gal).

### Colony Formation Assay

An equal number of cells (5,000 cells/well) were seeded in 6-well plates and cultured for ten days. Cells were then fixed with 1% paraformaldehyde/PBS and stained with 0.05% crystal violet. Wells were destained with 10% acetic acid. Absorbance (590nm) was measured using a spectrophotometer (Spectra Max 190).

### qRT-PCR

Total RNA was extracted from cells with Trizol and DNase treated, cleaned and concentrated using Zymo columns (Zymo Research). Optical density values of extracted RNA were measured using NanoDrop One (Thermo Scientific) to confirm an A260 and A280 ratios above 1.9. Relative expression of target genes listed in Table 1 were analyzed using the QuantStudio 3 Real-Time PCR System (Thermo Fisher Scientific) with clear 96 well plates (Greiner Bio-One). Primers were designed using the Integrated DNA Technologies (IDT) tool (https://www.idtdna.com/scitools/Applications/RealTimePCR/). Briefly, 5ng of total RNA was used to One-Step qPCR (Quanta BioSciences) following manufacturer’s instruction in a final volume of 10uL. Conditions for amplification were: 10 min at 48°C, 5 min at 95°C, 40 cycles of 10 s at 95°C and 7s at the corresponding annealing temperature. The assay ended with a melting-curve program: 15s at 95°C, 1 min at 70°C, then ramping to 95°C while continuously monitoring fluorescence. Each sample was assessed in triplicate. Relative quantification was determined normalizing to the reference gene *B2M* using the delta-delta Ct method.

### Chromatin Immunoprecipitation (ChIP)

ChIP was performed as previously described (42) using the ChIP-grade antibody mouse anti-H3K9me2 (Abcam). Cells were fixed in 1% paraformaldehyde for 5 min at room temperature and then quenched with 1 ml of 2.5 M glycine for 5 min at room temperature. Cells were washed twice with cold PBS. Cells were lysed in 1 ml ChIP lysis buffer (50 mM Hepes-KOH, pH 7.5, 140 mM NaCl, 1 mM EDTA, pH 8.0, 1% Triton X-100, and 0.1% deoxycholate [DOC] with 0.1 mM PMSF and the EDTA-free Protease Inhibitor Cocktail). Samples were incubated on ice for 10 min and then centrifuged at 3,000 rpm for 3 min at 4°C. The pellet was re- suspended in 500 μl lysis buffer 2 (10 mM Tris, pH 8.0, 200 mM NaCl, 1 mM EDTA, and 0.5 mM EGTA with 0.1 mM PMSF and the EDTA-free Protease Inhibitor Cocktail) and incubated at room temperature for 10 min. Samples were centrifuged at 3,000 rpm for 5 min at 4°C. Next, the pellet was resuspended in 300 μl lysis buffer 3 (10 mM Tris, pH 8.0, 100 mM NaCl, 1 mM EDTA, 0.5 mM EGTA, 0.1% DOC, and 0.5% *N*-lauroylsarcosine with 0.1 mM PMSF and the EDTA-free Protease Inhibitor Cocktail). Cells were sonicated using a Branson Sonifier 250 for four cycles of 10 seconds on 50 seconds off. Next, 30 μl of 10% Triton X-100 was added to each tube, and then samples were centrifuged at max speed for 15 min at 4°C. The supernatant was transferred to new tubes. 50 μl of the antibody bead conjugate solution was added, and chromatin was immunoprecipitated overnight on a rotator at 4°C. The following washes were performed for 15 min each by rotating for 15 min at 4°C: ChIP lysis buffer (twice), ChIP lysis buffer + 0.65 M NaCl, wash buffer (10 mM Tris-HCl, pH 8.0, 250 mM LiCl, 0.5% NP-30, 0.5% DOC, and 1 mM EDTA, pH 8.0), and TE (10 mM Tris-HCl, pH 8.0, and 1 mM EDTA, pH 8.0). DNA was eluted by incubating the beads with TES (50 mM Tris-HCl, pH 8.0, 10 mM EDTA, pH 8.0, and 1% SDS) for 30 min at 65°C. Reversal of cross-linking was performed by incubating samples overnight at 65°C. Proteins were digested using 1 mg/ml proteinase K and incubating at 37°C for 5 h. Finally, the DNA was purified using the Wizard SV Gel and PCR Clean Up kit (Promega). Immunoprecipitated DNA was analyzed by qPCR using iTaq Universal SYBR^®^ Green Supermix (BioRad). Conditions for amplification were: 5 min at 95°C, 40 cycles of 95°C for 10 sec and 30 sec with 62°C annealing temperature. The assay ended with a melting-curve program: 15s at 95°C, 1 min at 60°C, then ramping to 95°C while continuously monitoring fluorescence. Each sample was assessed in triplicate. Enrichment of H3K9me2 was determined by normalizing to a gene desert region (43). Primer sets used for ChIP-qPCR are detailed in Table 1.

### Western Blotting

Cells lysates were collected in 1X sample buffer (2% SDS, 10% glycerol, 0.01% bromophenol blue, 62.5mM Tris, pH 6.8, 0.1M DTT), boiled to 95°C for 10 min and sonicated. Protein concentration was determined using the Bradford assay. An equal amount of total protein was resolved using SDS-PAGE gels and transferred to nitrocellulose membranes (Fisher Scientific) at 110mA for 2 hours at 4°C. Membranes were blocked with 5% nonfat milk for 1 hour at room temperature. Membranes were incubated overnight at 4°C in primary antibodies in 4% BSA/TBS + 0.025% sodium azide. Membranes were washed 3 times in TBS-T for 5 min at room temperature after which they were incubated with HRP-conjugated secondary antibodies (Cell Signaling, Danvers, MA) for 1 hour at room temperature. After washing 3 times in TBS-T for 5 min at room temperature, proteins were visualized on film after incubation with SuperSignal West Pico PLUS Chemiluminescent Substrate (ThermoFisher, Waltham, MA).

### Quantification and Statistical Analysis

GraphPad Prism version 7.0 was used to perform statistical analysis. T-test and Oneway ANOVA followed by post hoc Tukey’s HSD tests were applied as appropriate. When indicated, P-values were adjusted according to Benjamini and Hochberg’s false discovery rate (FDR). The significance level was established at p < 0.05. Ns= not significant. Heatmaps were generated using the function *heatmap.2* from gplots R package v3.0-1. Minkowski distance and ward.D2 method of clustering were used to generate all heatmaps. Kaplan-Meier curves were generated using publicly available ovarian cancer mRNA gene chip data (44). Ovarian cancer patients were filtered by a serous histosubtype and *TP53* mutation to signify HGSC.

## Results

### Wildtype IDH1 is upregulated in high-grade serous ovarian cancer

To determine whether changes in metabolism correlate with ovarian cancer disease progression, we performed metabolomics on primary fallopian tube (FT) cells and HGSC cell lines. We observed a significant upregulation in the levels of all TCA cycle metabolites (**Fig. 1A**). However, we did not observe any consistent changes in glycolytic metabolites in HGSC cells (**Fig. S1A**). Indeed, metabolite profiling showed decreased lactate production in HGSC cells compared to FT cells (**Fig. S1B**). These data indicate that glucose is preferentially utilized by the TCA cycle in HGSC cells. To determine whether upregulation of enzyme expression is responsible for the observed increase in TCA cycle metabolism, we performed a RT-qPCR analysis of all 27 TCA cycle enzymes in five normal FT cell lines, including two primary and three immortalized lines, and four HGSC cell lines (**Fig. 1B**). All HGSC cell lines tested exhibited a significant increase in all TCA cycle enzyme expression except *IDH2* when compared to FT cells. Together, these data suggest that TCA cycle metabolism is universally altered in HGSC.

**Figure 1:**
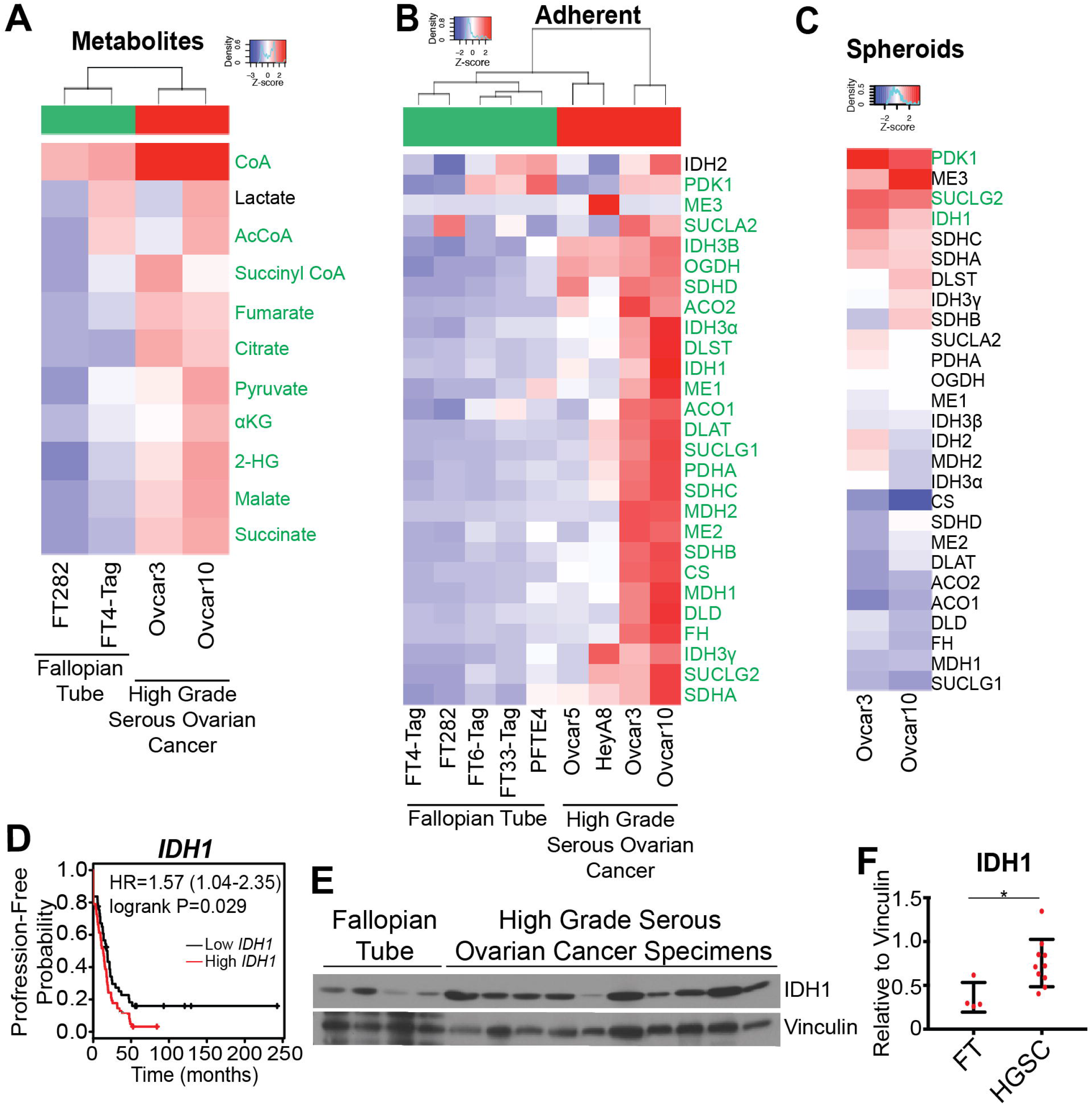
TCA cycle metabolism is upregulated in HGSC and increased expression of IDH1 correlates with decreased progression-free survival of HGSC patients. (A) LC/MS comparison of 2 fallopian tube (FT282 and FT4-Tag) and 2 HGSC (Ovcar3 and Ovcar10) cell lines. Heat map shows z-scores of relative abundance normalized to the mean of each individual TCA cycle metabolite. Significant metabolites are labeled in green. Adjusted p-value (FDR) <0.05. (B) Differentially expressed transcripts in 5 parental fallopian tube (FT282, FT4-Tag, FT6-Tag, FT33-Tag, and PFTE4) and 4 HGSC (HeyA8, Ovcar5, Ovcar3, and Ovcar10) cell lines. Heat map shows z-scores of expression levels relative to *B2M* of 27 TCA cycle enzymes. Significant transcripts are labeled in green. Adjusted p-value (FDR) <0.05 (C) Differentially expressed transcripts of 27 TCA cycle enzymes. Heat map shows fold change z-scores of indicated HGSC cell lines cultured in ultra-low attachment and adherent conditions. Significant transcripts are labeled in green. Adjusted p-value (FDR) <0.001 (D) Kaplan-Meier curve for disease progression-free survival of ovarian cancer patients classified according to *IDH1* expression level. Ovarian cancer patients were filtered for a serous histosubtype and *TP53* mutation. Estimates of the hazard ration (HR) and logrank p-values are indicated. (E) Immunoblot analysis of IDH1 comparing normal human fallopian tube and HGSC patient tissue lysates. Vinculin was used as a loading control. (F) IDH1 protein expression normalized to vinculin and quantified using ImageJ. Data represent mean ± SD. *p<0.03.

HGSC disseminates to the peritoneal cavity in the form of spheroids during late stage disease (45, 46). In order to mimic these conditions *in vitro*, ultra-low attachment (ULA) plates were used to induce the formation of spheroids. As Ovcar3 and Ovcar10 cells showed a robust increase in TCA cycle metabolites and TCA cycle enzymes (**Fig. 1A-B**), we compared TCA cycle enzyme expression in these cells from adherent and spheroid conditions (**Fig. 1C**). Under these conditions, only three TCA cycle enzymes showed increased expression: *PDK1* (pyruvate dehydrogenase kinase 1), *SUCLG2* (succinate-CoA ligase GDP-forming beta subunit), and *IDH1* (p<0.001). Further analysis of HGSC patients demonstrated that only high *IDH1* expression was associated with a significant decrease in progression-free survival (**Fig. 1D** and S1C-D). Consistently, wildtype IDH1 protein is increased in primary HGSC patient tumors when compared to normal FT tissues (**Fig. 1E-F**). Together, these data demonstrate that wildtype IDH1 is significantly upregulated in HGSC compared to FT.

### Inhibition and knockdown of wildtype IDH1 inhibits proliferation of HGSC cells

Next, we sought to determine whether upregulation of IDH1 is critical for HGSC proliferation. Therefore, we examined the effects of pharmacological inhibition and knockdown of IDH1 on HGSC cell proliferation. There are currently no commercially available inhibitors of wildtype IDH1; therefore, we used an IDH1 mutant inhibitor (IDH1i) that also targets the wildtype protein at higher concentrations (30, 47). Under adherent conditions, inhibition of IDH1 in Ovcar3 cells decreased BrdU incorporation and colony formation (**Fig. 2A-D**). Similar results were observed in Ovcar10 cells, indicating this is not a cell line-specific phenomenon (**Fig. S2A-D**). We examined genetic knockdown of IDH1 in Ovcar3 cells using two independent hairpins (**Fig. 2E**). Similar to the IDH1 inhibitor, knockdown of IDH1 decreased BrdU incorporation, colony formation, and cyclin A expression (**Fig. 2F-J**). Similar results were observed upon IDH1 knockdown in Ovcar10 cells (**Fig. S2E-I**). Similar to adherent conditions, proliferation was also decreased upon IDH1 inhibition in spheroid conditions as indicated by decreased cyclin A (**Fig. 2K**). Taken together, we conclude that inhibition or knockdown of wildtype IDH1 suppresses proliferation of HGSC cells.

**Figure 2:**
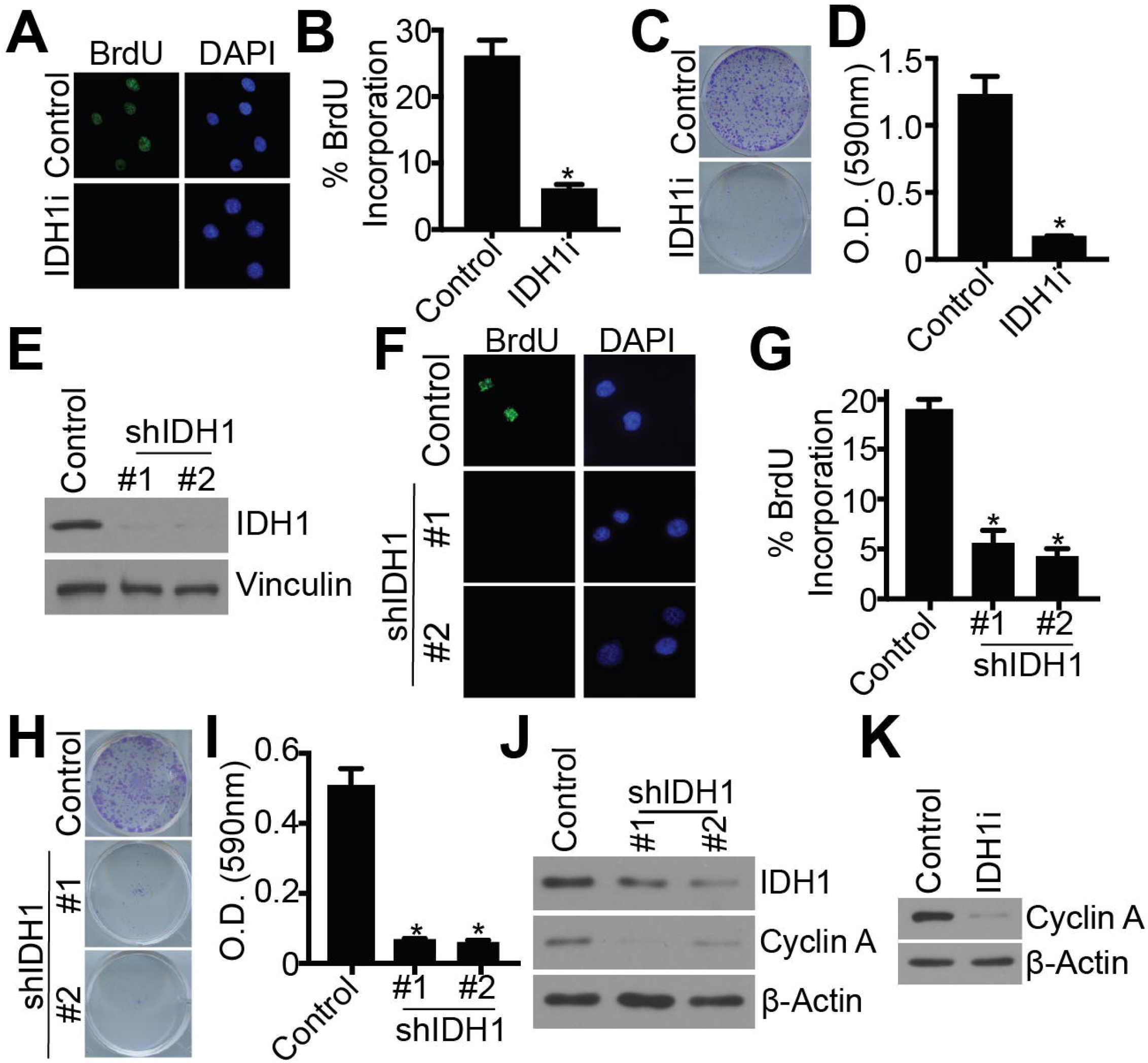
Pharmacological inhibition and knockdown of IDH1 suppresses proliferation of Ovcar3 HGSC cells. (A-D) Ovcar3 cells were treated with either DMSO or 15 μM GSK864 (IDH1i) for 7 days. (A) BrdU incorporation. One of three experiments shown. (B) Quantification of (A). Data represent mean ± SD. *p<0.0001 (C) Colony formation. Cells were seeded at equal densities and 10 days later stained with 0.05% crystal violet. One of three experiments is shown. (D) Quantification of (C). Data represent mean ± SD. *p<0.0001 (E-J) Ovcar3 cells were infected with lentivirus expressing two independent short hairpin RNAs (shRNAs) targeting IDH1 (shIDH1). Empty vector was used as a control. Cells were selected with puromycin for 7 days. (E) Immunoblot analysis of IDH1. Vinculin was used as a loading control. One of 5 experiments is shown. (F) BrdU incorporation. One of three experiments shown. (G) Quantification of (F). Data represent mean ± SD. *p<0.0001 (H) Colony formation. Cells were seeded at equal densities and 10 days later stained with 0.05% crystal violet. One of three experiments is shown. (I) Quantification of (H). Data represent mean ± SD. *p<0.0001 (J) Immunoblot analysis of IDH1 and cyclin A. β-Actin was used as a loading control. One of three experiments shown. (K) Immunoblot of Ovcar3 cells cultured in ULA conditions and treated with 15 μM GSK864 (IDH1i) for cyclin A. β-Actin was used as a loading control. One of three experiments shown.

### Inhibition and knockdown of IDH1 induces senescence of HGSC cells

Next, we sought to determine the mechanism by which inhibition and knockdown of IDH1 suppresses HGSC cell proliferation. In adherent conditions, knockdown of IDH1 did not induce cell death (Fig S3A). Interestingly, upon suppression of IDH1, cells exhibited a large and flat morphology, which are characteristic of senescence (**Fig. S3B**) (48); therefore, we investigated whether suppression of proliferation was due to senescence induction. Towards this goal, we examined the expression of senescence-associated beta galactosidase (SA-β-Gal) activity. Indeed, inhibition of IDH1 increased SA-β-Gal activity when compared to control Ovcar3 cells (**Fig. 3A-B**). Other senescent markers such as decreased lamin B1 and increased PML bodies were also observed when IDH1 was inhibited (**Fig. 3C-E**). Similar results were observed in IDH1 knockdown cells (**Fig. 3F-J**). However, the senescence-associated secretory phenotype (SASP) (49) was not robustly increased in IDH1 knockdown cells when compared to etoposide treated cells (**Fig. S3C**). Senescence was also observed in Ovcar10 cells upon IDH1 inhibition or knockdown, suggesting this is not a cell line specific effect (**Fig. S3D-L**). Additionally, knockdown or inhibition of IDH1 induced senescence in spheroid conditions as indicated by decreased lamin B1 (**Fig. 3K** and **Fig. S3M**). IDH1 produces αKG in a reversible reaction (50). Addback of exogenous αKG in IDH1 knockdown partially rescued the senescent phenotype (**Fig 3L-O**), suggesting HGSC preferentially utilizes the forward oxidative decarboxylation reaction to produce αKG. These data suggest that inhibition or knockdown of IDH1 induces senescence in both adherent and spheroid conditions and may allow for a sustained anti-tumor response due to a lack of SASP gene expression.

**Figure 3:**
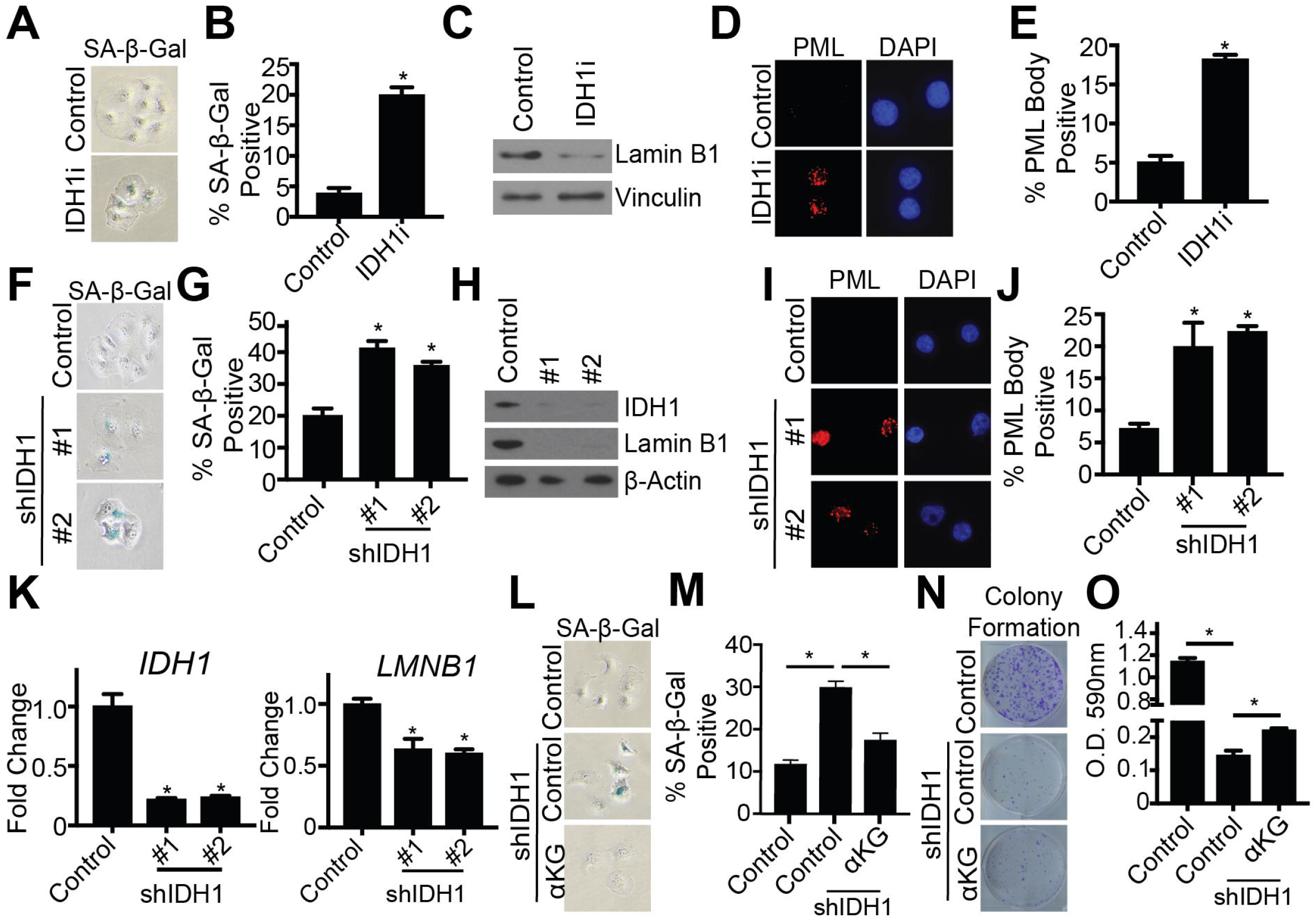
Inhibition and knockdown of IDH1 induces senescence in HGSC in both adherent and spheroid conditions. (A-E) Ovcar3 cells were treated with either DMSO or 15μM GSK864 (IDH1i) for 7 days. (A) SA-β-Gal activity. One of three experiments is shown. (B) Quantification of (A). Data represent mean ± SD. *p<0.0001 (C) Immunoblot analysis of lamin B1. One of three experiments is shown. (D) PML body foci. One of three experiments is shown. (E) Quantification of (D). Data represent mean ± SD. *p<0.0001 (F-J) Ovcar3 cells were infected with lentivirus expressing two independent shRNAs targeting IDH1. Empty vector was used as a control. Cells were selected in 3μg/mL puromycin for 7 days. (F) SA-β-Gal activity. One of three experiments is shown. (G) Quantification of (F). Data represent mean ± SD. *p<0.0001 (H) Immunoblot analysis of IDH1 and lamin B1. One of three experiments is shown. (I) PML body foci. One of three experiments is shown. (J) Quantification of (I). Data represent mean ± SD. *p<0.0005 (K) qRT-PCR analysis of *IDH1* and *LMNB1* (encoding Lamin B1) in Ovcar3 shIDH1 ULA. RNA was collected four days after infection. Data represent mean ± SD. *p<0.0002 (L-O) Ovcar3 cells were infected with lentivirus expressing one shRNA targeting IDH1 (#1). Empty vector was used as a control. Cells were selected in 3μg/mL puromycin for 7 days. Exogenous αKG (1mM) was added to the indicated cells. (L) SA-β-Gal activity of shIDH1 with and without αKG. One of three experiments is shown. (M)Quantification of (L). Data represent mean ± SD. *p<0.0001 (N) Colony formation of shIDH1 with and without αKG. One of three experiments is shown. (O) Quantification of (N). Data represent mean ± SD. *p<0.002

### Knockdown of IDH1 induces senescence through increased repressive histone methylation at proliferation-promoting gene loci

Next, we aimed to determine the mechanism by which targeting IDH1 induces senescence. Inhibition of IDH1 has previously been shown to increase reactive oxygen species (51), which is a known inducer of senescence (52). However, we did not observe an increase in ROS upon IDH1 knockdown (**Fig. S4A**), and treatment with the antioxidant N-acetyl cysteine (NAC) did not rescue senescence due to IDH1 knockdown (**Fig. S4B**). Moreover, we did not observe an increase in DNA damage, another well-known mechanism of therapy-induced senescence (**Fig. S4C**). Increased repressive histone methylation is observed in senescence. In particular, induction of senescence is due in part to increased histone H3 lysine 9 di-methylation (H3K9me2) at proliferation-promoting gene loci (13, 14, 53). Increased histone methylation may be due to decreased histone demethylase activity, and αKG, the product of IDH1 activity, acts as a co-factor histone demethylases (31, 32). Therefore, we investigated whether knockdown of IDH1 modulates repressive histone methylation of proliferation-promoting genes to induce senescence. Chromatin-immunoprecipitation (ChIP) assays using an antibody specific for repressive H3K9 dimethylation (H3K9me2) demonstrated increased H3K9me2 at *PCNA* and *MCM3* gene loci in both Ovcar3 and Ovcar10 cells with IDH1 knockdown compared to controls (**Fig. 4A**). Consistently, *PCNA* and *MCM3* mRNA expression was decreased in these cells (**Fig. 4B**). Together, these data suggest that knockdown of IDH1 increases histone methylation at proliferation-promoting gene loci to decrease their expression and induce senescence.

**Figure 4:**
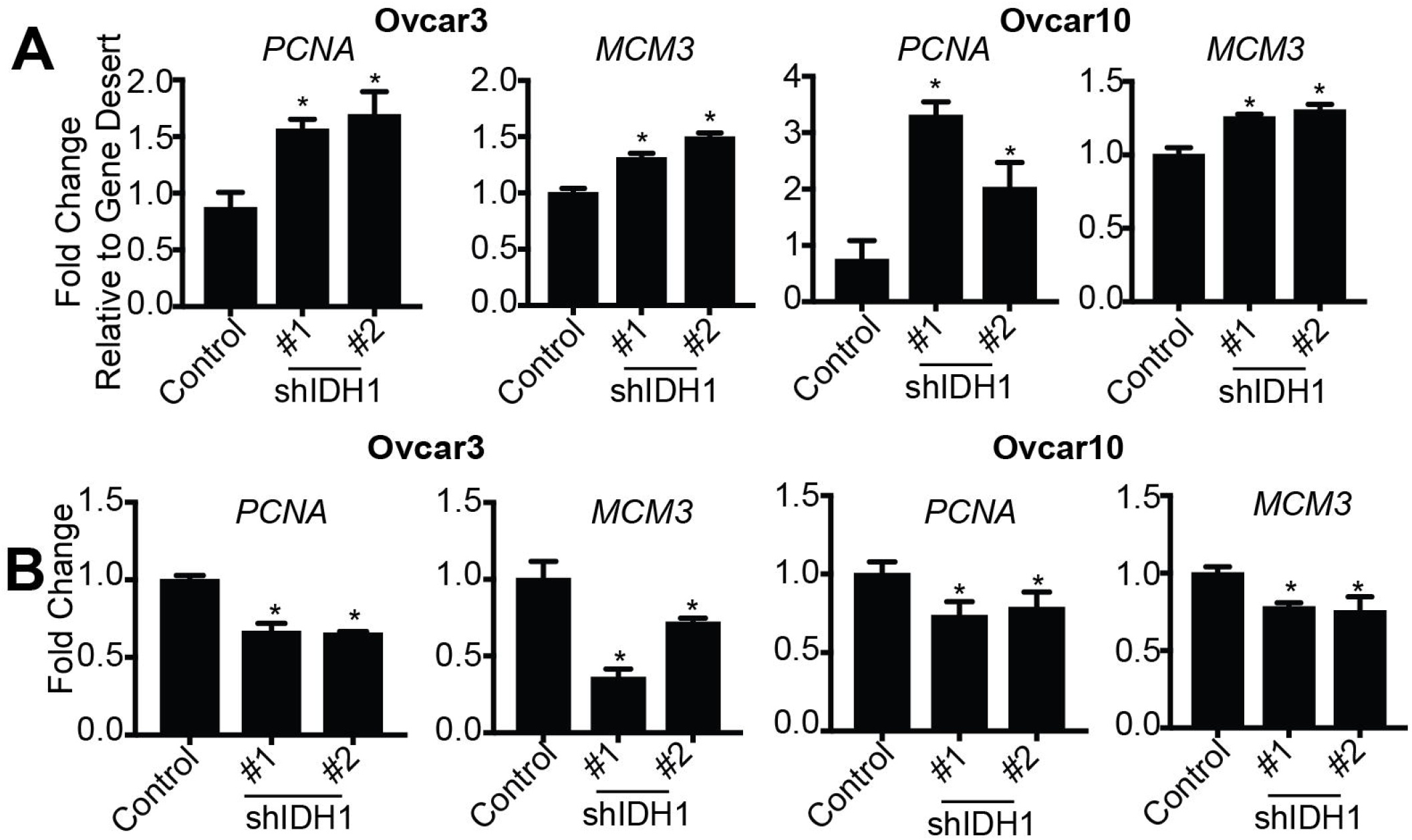
Knockdown of IDH1 increases repressive H3K9me2 at proliferation-promoting gene loci. (A&B) Ovcar3 and Ovcar10 cells were infected with two independent hairpins targeting IDH1. Empty vector was used as a control. Cells were selected with puromycin for 7 days. (A) Chromatin immunoprecipitation qPCR analysis of H3K9me2 at the loci of *PCNA* and *MCM3*. Data is normalized to a gene desert region. One of two experiments is shown. Data represent mean ± SD. *p<0.02 (B) RT-qPCR analysis *PCNA* and *MCM3* expression. *B2M* was used as a reference gene. One of three experiments is shown. Data represent mean ± SD. *p<0.008

## Discussion

In this study, we found that the TCA cycle enzyme IDH1 is significantly increased in HGSC cells compared to FT cells. Inhibition or knockdown of IDH1 suppressed HGSC cell proliferation and induced senescence, a stable cell cycle arrest. Mechanistically, targeting IDH1 induced senescence by increasing repressive H3K9me2 at the loci of proliferation-promoting genes, *PCNA* and *MCM3*. Patient HGSC samples indicated a significant increase in IDH1 when compared to normal FT, which correlated with worse progression-free survival. Taken together, these data indicate that high expression of IDH1 is a poor prognostic factor for HGSC and inhibiting its activity represents a novel strategy for HGSC progression and dissemination.

Approximately 70% of HGSC receiving platinum-based standard-of-care therapy will relapse with chemoresistant disease (4, 54). Currently, there are few therapies available for these patients after relapse, ultimately leading to patient mortality. Aberrant metabolism is a global characteristic of cancer cells (20), and altered metabolism of cancer cells affects the response to many chemotherapies (55–57). Therefore, targeting metabolism may serve as a novel therapeutic in many cancer types. Previous studies have implicated altered metabolic pathways in ovarian cancer (58, 59), and multiple studies have targeted glycolytic enzymes for ovarian cancer treatment (60–62). However, our study demonstrates that FT cells also undergo glycolysis (**Fig. S1A-B**), suggesting that targeting this pathway may result in toxicity to normal tissue. We found that TCA cycle metabolites are significantly upregulated in HGSC cells compared to FT, which suggests that targeting this pathway may lead to less toxicity. Specifically, we identified wildtype IDH1 as a potential target for HGSC. Recently, the IDH1 inhibitor Tibsovo (ivosidenib, AG-120) was approved for relapsed AML with mutant IDH1 (63). This inhibitor is also effective against wildtype IDH1, but not wildtype IDH2 (64). Our data and others suggest that targeting wildtype IDH1 is a rational therapy for multiple cancers (65).

We found that targeting wildtype IDH1 induces a sustained proliferative arrest in HGSC cell lines through induction of senescence. While IDH1 has never been explored in ovarian cancer, this is consistent with previous publications in glioblastoma (30, 66). This study found that silencing IDH1 expression in glioblastoma reduces NADPH to induce senescence. NADPH is critical for multiple cellular processes, including nucleotide synthesis and redox homeostasis (67). Indeed, they found that senescence was rescued by supplementation with both nucleotides and the antioxidant N-acetyl-L-cysteine. Consistently, other groups have shown that reductive carboxylation of IDH1 is increased under anchorage-independent conditions to mitigate reactive oxygen species (ROS) (68). While these studies linked IDH1 to increased oxidative stress, we did not observe a significant increase in reactive oxygen species in HGSC cells with IDH1 knockdown (Fig S4A-B). This suggests that there is a context or cell type-dependent regulation of senescence by targeting IDH1.

Late stage HGSC primarily disseminates to the peritoneal cavity in the form of spheroids (69, 70). These cells then seed on organs within the peritoneal cavity, which in part leads to the morbidity and mortality of late-stage ovarian cancer (1). We found that inhibition and shRNA-mediated knockdown of IDH1 induced senescence in both adherent and spheroid conditions (**Fig. 3**). This observation suggests that targeting IDH1 may be a therapeutic strategy for HGSCs at various stages including early and late stage diagnoses. Moreover, HGSCs are universally characterized by p53 mutations (71–73) and ~20% have decreased *RB1* (encoding for retinoblastoma protein, RB) (74). While these two pathways are implicated in the senescence-associated cell cycle arrest (75, 76), our data suggest that targeting IDH1 downregulates proliferation promoting genes through epigenetic modifications independent of p53 and RB status.

Senescence is characterized by marked chromatin remodeling and increased repressive histone methylation, collectively termed the senescence-associated heterochromatin foci (SAHF) (13, 14). Specifically, di- and tri-methylation of histone 3 lysine 9 at the loci of proliferation promoting genes is implicated in senescence. We found that knockdown of IDH1 induced H3K9 methylation at *PCNA* and *MCM3* gene loci (**Fig. 4A**), two E2F1-responsive proliferation-promoting genes involved in senescence induction (13). This led to a decrease in expression of these genes (**Fig. 4B**). Previous publications found that mutant IDH1, through the oncometabolite 2-hydroxyglutarate (2HG), alters histone methylation at DNA damage response and differentiation gene loci through competitive inhibition of histone demethylases that require αKG for their activity (77, 78). Alterations in histone demethylases have been connected to senescence induction (36, 79, 80). Our data suggest that demethylated H3K9 is critical for ovarian cancer cells to proliferate, and targeting this pathway is a novel therapeutic to inhibit ovarian cancer proliferation via senescence. Future work is needed to determine the specific histone H3K9 demethylase responsible for the senescent phenotype.

Senescence is considered a tumor suppressive mechanism, and senescence induction is considered a beneficial therapeutic outcome (16, 19, 81). However, senescence may also promote cancer through the senescence-associated secretory phenotype (SASP), which increases inflammation and alters the surrounding microenvironment milieu (82). We found that SASP gene expression was not increased upon IDH1 knockdown (**Fig. S3C**), suggesting that targeting IDH1 may lead to a sustained cell cycle arrest and therapeutic response. Our results in addition to others suggest that senescence induction in the cell-type specific context of ovarian cancer may overall be tumor-suppressing (17, 19, 83). Altogether, we propose that targeting IDH1 may act as a novel pro-senescent therapy for HGSC patients.

## Authors’ Contributions

Conception and design: K.M. Aird and E.S. Dahl Development of methodology: N.W. Snyder

Acquisition of data (acquired and managed patients, provided facilities, etc.): E.S. Dahl, R. Buj, K.E. Leon, J. M. Newell, B.G. Bitler, N.W. Snyder, K.M. Aird

Analysis and interpretation of data (e.g., statistical analysis, biostatistics, computational analysis): R. Buj and E.S. Dahl

Writing, review, and/or revision of the manuscript: E.S. Dahl, B. G. Bitler, N.W. Snyder, and K.M. Aird

Study supervision: B.G. Bitler, K.M. Aird and N.W. Snyder

## Acknowledgments

This work was partially supported by grants from the National Institutes of Health (R00CA194309 to K.M.A., R00CA194318 to B.G.B, R03CA211820 to N.W.S, and the use of core facilities under P30ES013508) and the W. W. Charitable Trust (to K.M.A.).

We would like to thank the members of the Aird Lab for their thoughtful contributions. We would like to thank Dr. Ronny Drapkin (University of Pennsylvania) and Dr. Anna Loshkin (University of Pittsburgh) for the fallopian tube cells.

## Supplemental Information

### Supplemental Materials and Methods

#### N-acetyl-L-cysteine supplementation

Cells were treated with 500 μM n-acetyl cysteine (NAC Sigma Aldrich) where indicated.

#### YSI Metabolite Measurement

Lactate production was measured using a YSI 2950 Bioanalyzer (Yellow Springs, OH). Briefly, the same number of cells were seeded in 12-well plates and media was changed 24-hours later. The next day, fresh media was harvested and cells were counted to normalize for proliferation.

#### Flow Cytometry

7AAD: Cells were trypsinized and washed once with 1X PBS. Cells were then resuspended and 5μL of 7-aminoactinomycin D (7AAD) (Tonbo Biosciences) was added. Live cells were run through a 10-color FACSCanto flow cytometer. DHE: HGSC cells were incubated for 30 min with 10μM Dihydroethidium (DHE abcam). Cells were then washed and run through a 10-color FACSCanto flow cytometer. Positive control cells were incubated with 100μM H_2_O_2_ prior to DHE treatment. Data was analyzed using FlowJo software (Ashland, OR).

**Supplemental Figure 1:**
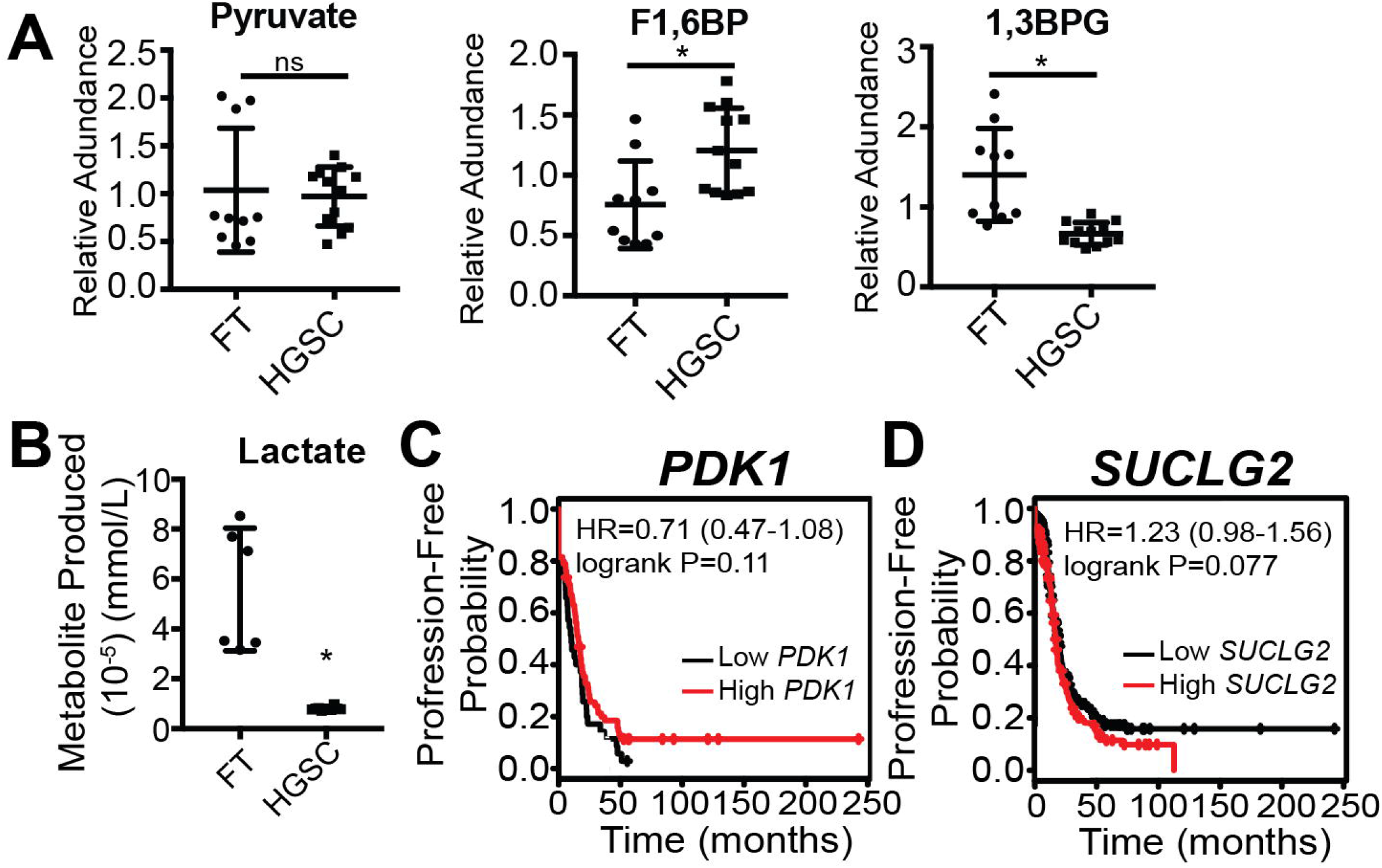
Glycolytic metabolites are not consistently altered in HGSC compared to fallopian tube cells; increased expression of TCA cycle enzymes *PDK1* and *SUCLG2* do not correlate with decreased progression-free survival. Related to Figure 1. (A) LC/MS comparison of fallopian tube cell lines (FT282 and FT4-Tag) and HGSC cell lines (Ovcar3 and Ovcar10). Relative abundance of indicated metabolites is shown. F1,6BP (fructose 1,6-bisphosphate) and 1,3BPG (1,3-bisphosphoglyceric acid). Data represent mean ± SD. *p<0.009. (B) Lactate production of FT and HGSC cells utilizing a YSI-bioanalyzer. Data was normalized to control media and cell number. Data represent mean ± SD. *p<0.0008. (C) Kaplan-Meier curve for disease progression-free survival of ovarian cancer patients classified according to *PDK1* expression level. Ovarian cancer patients were filtered for a serous histosubtype and *TP53* mutation. Estimates of the hazard ration (HR) and logrank p-values are indicated. (D) Kaplan-Meier curve for disease progression-free survival of ovarian cancer patients classified according to *SUCLG2* expression level. Ovarian cancer patients were filtered for a serous histosubtype and *TP53* mutation. Estimates of the hazard ration (HR) and logrank p-values are indicated.

**Supplemental Figure 2:**
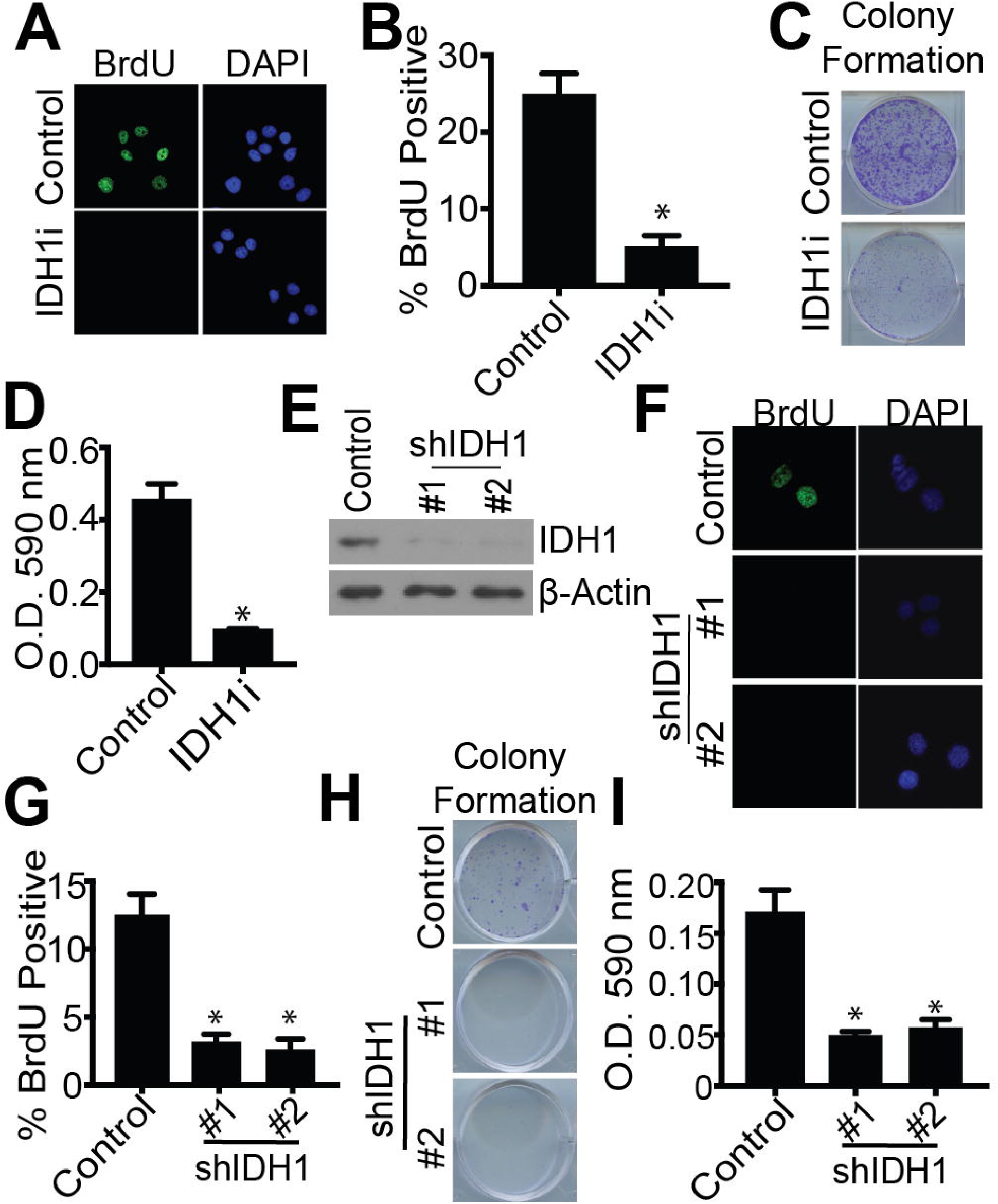
Pharmacological inhibition and knockdown of IDH1 suppresses proliferation of Ovcar10 HGSC cells. Related to Figure 2. (A-D) Ovcar10 cells were treated with either DMSO or 15μM GSK864 (IDH1i) for 7 days. (A) BrdU incorporation. One of three experiments shown. (B) Quantification of (A). Data represent mean ± SD. *p<0.0001 (C) Colony formation. Cells were seeded at equal densities and 10 days later stained with 0.05% crystal violet. One of three experiments is shown. (D) Quantification of (C). Data represent mean ± SD. *p<0.0001 (E-I) Ovcar10 cells were infected with lentivirus expressing two independent short hairpin RNAs (shRNAs) targeting IDH1 (shIDH1). Empty vector was used as a control. Cells were selected with puromycin for 7 days. (E) Immunoblot analysis of IDH1. β-Actin was used as a loading control. One of 5 experiments is shown. (F) BrdU incorporation. One of three experiments shown. (G) Quantification of (G). Data represent mean ± SD. *p<0.0001 (H) Colony formation. Cells were seeded at equal densities and 10 days later stained with 0.05% crystal violet. One of three experiments is shown. (I) Quantification of (H). Data represent mean ± SD. *p<0.0001

**Supplemental Figure 3:**
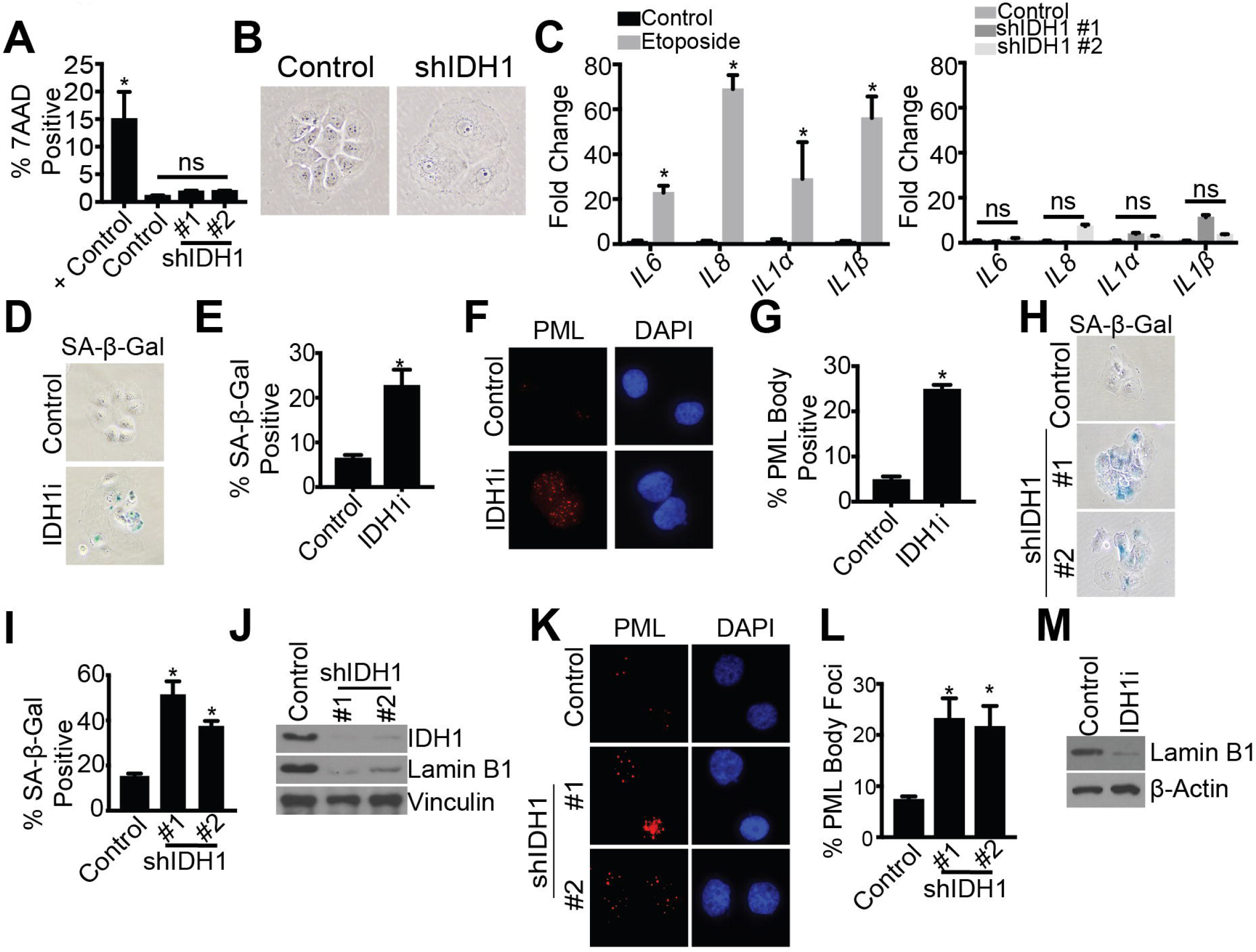
Knockdown of IDH1 induces senescence in Ovcar10 cells but does not increase SASP gene expression. Related to Figure 3. (A&B) Ovcar10 cells were infected with lentivirus hairpins two independent shRNAs targeting IDH1. Empty vector was used as a control. Cells were selected in puromycin for 7 days. (A) 7AAD flow cytometry analysis. One of three experiments is shown. *p<0.0005 (B) Brightfield image of cell morphology. (C) Ovcar10 cells were infected with lentivirus hairpins two independent shRNAs targeting IDH1. Empty vector was used as a control. Cells were selected in puromycin for 7 days. qRT-PCR analysis of SASP gene expression *(IL6, IL8, IL1α, IL1β)*. Etoposide (10μM) was used as a positive control. One of three experiments is shown. Data represent mean ± SD. *p<0.0001 (D-G) Ovcar10 cells were treated with either DMSO or 15μM GSK864 (IDH1i) for 7 days. (D) SA-β-Gal activity. One of three experiments is shown. (E) Quantification of (D). Data represent mean ± SD. *p<0.002. (F) PML body foci. One of three experiments is shown. (G) Quantification of (F). Data represent mean ± SD. *p<0.0001 (H-L) Ovcar10 cells were infected with lentivirus hairpins two independent shRNAs targeting IDH1. Empty vector was used as a control. Cells were selected in puromycin for 7 days. (H) SA-β-Gal activity. One of five experiments is shown. (I) Quantification of (H). Data represent mean ± SD. *p<0.0001 (J) Immunoblot analysis of IDH1 and lamin B1. One of three experiments is shown. (K) PML body foci. One of three experiments is shown. (IDH1i) for 4 days. Immunoblot analysis of lamin B1. One of two experiments is shown.

**Supplemental Figure 4:**
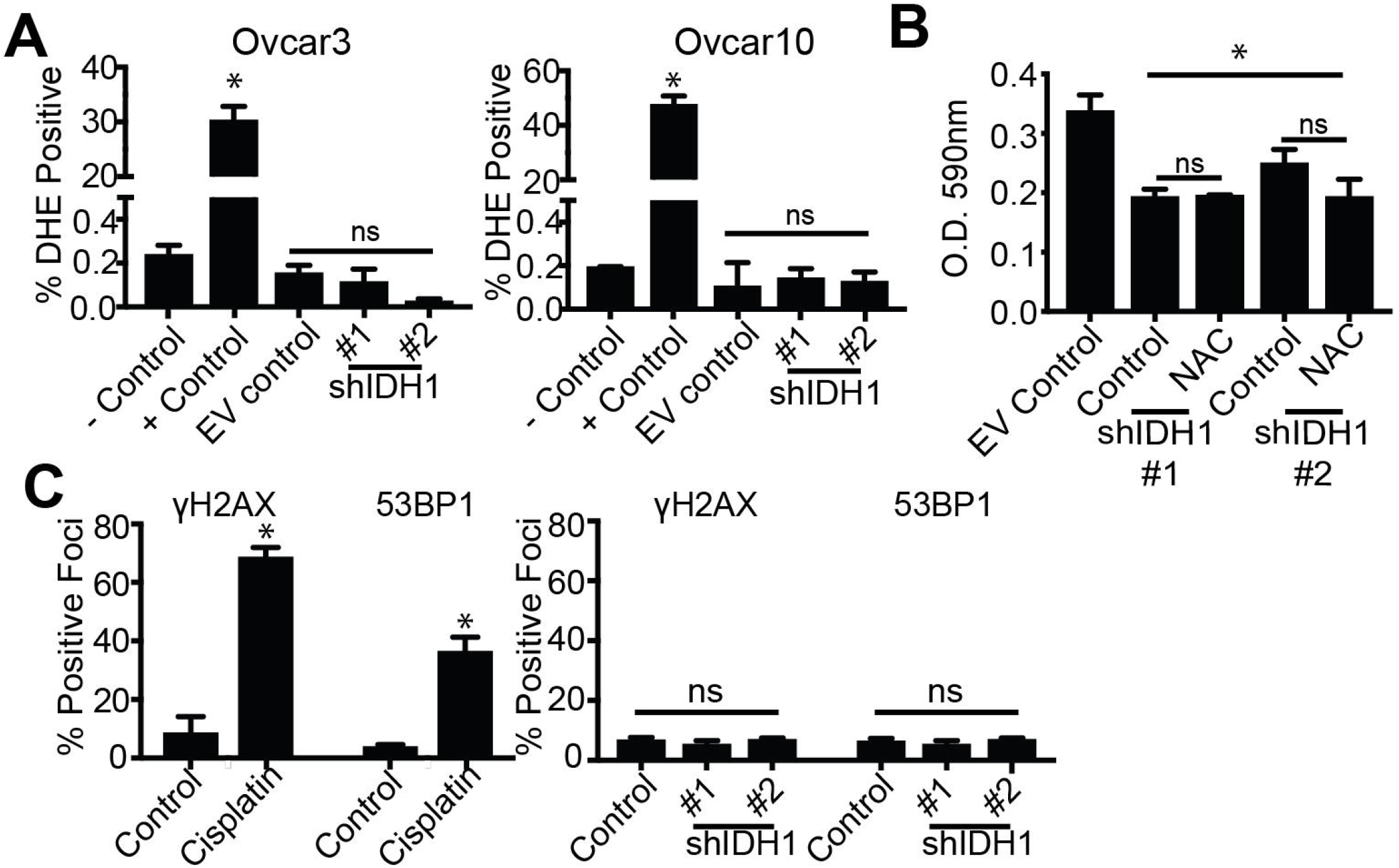
Senescence induced by IDH1 knockdown is independent of ROS and DNA damage. Related to Figure 4. (A) Ovcar3 and Ovcar10 cells were infected with two independent hairpins targeting IDH1. Cells were selected with puromycin for 7 days. Flow cytometry analysis of ROS using DHE as a marker. Cisplatin was used as a positive control and DMF was used as a negative control. Empty vector (EV) was also used as a negative control. One of three experiments is shown. Data represent mean ± SD. *p<0.0001 (B) Ovcar10 cells were infected with two independent hairpins targeting IDH1. Cells were selected with puromycin for 7 days. 500μM of n-acetyl cysteine (NAC) or vehicle control (water) was added to cells. Cells were seeded in 6-well plates and stained with crystal violet. Quantification O.D. 590 was determined. One of two experiments is shown. Data represent mean ± SD. *p<0.005 (C) Ovcar3 cells were infected with two independent hairpins targeting IDH1. Cells were selected with puromycin for 7 days. Immunofluorescence quantification of 53BP1 and γH2AX. Cisplatin (1μM) was used as a positive control. One of three experiments is shown. Data represent mean ± SD. *p<0.02.

